# Transcriptome-based screening in TARDBP/TDP-43 knock-in motor neurons identifies the NEDD8-activating enzyme inhibitor MLN4924

**DOI:** 10.1101/2024.01.30.578100

**Authors:** Sarah Lépine, Gilles Maussion, Alexandria Schneider, Angela Nauleau-Javaudin, María José Castellanos-Montiel, Georgina Jiménez Ambriz, Dan Spiegelman, Narges Abdian, Anna Krystina Franco-Flores, Ghazal Haghi, Lale Gursu, Mathilde Chaineau, Thomas Martin Durcan

**Author notes:** Corresponding authors: Thomas M. Durcan, 3801 University Street, Montreal, QC, H3A 2B4, Canada; Mathilde Chaineau, 3801 University Street, Montreal, QC, H3A 2B4, Canada. **Conflict-of-interest statement** The authors have declared that no conflict of interest exists.

## Abstract

A growing body of knowledge implicates perturbed RNA homeostasis in amyotrophic lateral sclerosis (ALS), a neurodegenerative disease that currently has no cure and few available treatments. Dysregulation of the multifunctional RNA-binding protein TDP-43 is increasingly regarded as a convergent feature of this disease, evidenced at the neuropathological level by the detection of TDP-43 pathology in most patient tissues, and at the genetic level by the identification of disease-associated mutations in its coding gene *TARDBP*. To characterize the transcriptional landscape induced by *TARDBP* mutations, we performed whole-transcriptome profiling of motor neurons differentiated from two knock-in iPSC lines expressing the ALS-linked TDP-43 variants p.A382T or p.G348C. Our results show that the *TARDBP* mutations significantly altered the expression profiles of mRNAs and microRNAs of the 14q32 cluster in MNs. Using mutation-induced gene signatures and the Connectivity Map database, we identified compounds predicted to restore gene expression toward wild-type levels. Among top-scoring compounds selected for further investigation, the NEDD8-activating enzyme inhibitor MLN4924 effectively improved cell viability and neuronal activity, highlighting a possible role for protein post-translational modification via NEDDylation in the pathobiology of TDP-43 in ALS.

## Introduction

Amyotrophic lateral sclerosis (ALS) is a neurodegenerative disorder caused by the progressive degeneration of motor neurons (MNs) in the brain and the spinal cord resulting in weakness, loss of ambulation, and eventual fatal paralysis of respiratory function.^1^ From disease onset, the mean survival ranges from two to four years.^2^ At present, the few available treatments (i.e., riluzole,^3–5^ edaravone,^6,7^ AMX0035^8–10^) offer modest benefits or are only efficacious in a subset of patients, as for the SOD1-lowering therapy tofersen.^11–13^ Our limitations in treating these patients reflect our incomplete understanding of the molecular basis of ALS and the difficulty in therapeutically addressing the multifactorial nature of the disease.

Several lines of evidence implicate disturbed RNA metabolism in ALS. A pathological hallmark of ALS is the nuclear depletion and cytoplasmic aggregation of TAR DNA-binding protein 43 (TDP-43), an RNA-binding protein encoded by *TARDBP* involved in nearly all aspects of RNA metabolism.^14–16^ Mutations in *TARDBP* and several other genes encoding RNA-binding proteins (e.g., *FUS*, *HNRNPA1*, *HNRNPB2*, *MATR3, TAF15*) have been implicated in ALS (ALSoD database; https://alsod.ac.uk/). Furthermore, transcriptome alterations have been repeatedly reported in the brain and spinal cord of patients who have succumbed to this disease,^17–22^ establishing a strong link between perturbed RNA homeostasis and ALS.

Initially identified as a transcriptional repressor,^23^ TDP-43 has since been implicated in splicing,^24^ microRNA (miRNA) biogenesis,^25,26^ RNA stability and transport,^27,28^ and translation.^29^ In keeping with these functions, TDP-43 localizes to ribonucleoprotein (RNP) condensates both in the nucleus (i.e., nuclear speckles,^30^ paraspleckles,^31^ and Cajal bodies^31^) and the cytoplasm (i.e., transport granules,^27,28^ stress granules,^32–34^ and processing (P)-bodies^29^), which broadly serve to regulate gene expression in time and space. While the many roles of TDP-43 in RNA metabolism have been extensively documented, the impact of disease-associated mutations in *TARDBP* on the RNA landscape is still being determined. Several studies have reported RNA abnormalities in *TARDBP*/*Tardbp* mutant mouse models,^35–40^ but whether the observed changes accurately reflect the human condition has been questioned.

As dysregulation of gene expression can have broad downstream consequences, determining the transcriptome alterations that arise in ALS can inform the development of transcriptome-correcting therapies. This drug discovery paradigm, known as “transcriptome reversal”,^41^ could give rise to therapies able to normalize several disease pathways simultaneously.

To this end, we performed whole-transcriptome profiling of motor neurons differentiated from two knock-in iPSC lines expressing the ALS-linked TDP-43 variants p.A382T or p.G348C, leading to the identification of mutation-induced RNA signatures. Positing that shared alterations may be reflective of underlying disease mechanisms, we queried the Connectivity Map (CMap) database^42^ for compounds predicted to normalize gene expression changes toward wild-type levels. Selected top-scoring compounds identified *in silico* were tested experimentally for their ability to ameliorate previously identified disease-relevant readouts^43^ in two phenotypic screens for viability and neuronal activity. This approach led to the identification of the NEDD8-activating enzyme inhibitor MLN4924.

## Results

### Transcriptomic profiling of MNs differentiated from *TARDBP* knock-in iPSCs

We have recently reported the generation of two homozygous knock-in iPSC lines with mutations in *TARDBP* coding for TDP-43^A382T^ or TDP-43^G348C^, two frequent ALS variants of TDP-43.^43^ To characterize the transcriptomic profiles of MNs differentiated from those cells, we performed next-generation RNA sequencing (RNA-seq) on total RNA extracted from mutant and isogenic control iPSC-derived MN cultures at 4-weeks post-plating (**Figure 1A** and **Supplemental Table S1**). Analysis of normalized counts confirmed the expression of the MN markers *ISL1*, *MNX1* (Hb9), *CHAT*, and *SLC18A3* (VAChT) (**Supplemental Figure 1A**). Unsurprisingly, *MNX1* (Hb9) was detected at relatively low levels as this early-MN marker is gradually downregulated with maturation in limb-innervating MNs.^44^ In addition, we noted the expression of the V2 interneuron markers *LMO4*, *VSX2*/*CHX10*, and *SOX14*, reflecting the anatomical proximity of V2 and pMN domains in the developing spinal cord. Markers for astrocytes, oligodendrocytes, neuronal progenitors, and V1/V3 interneurons were mostly absent or expressed at low levels (**Supplemental Figure 1A**). Expression of the stem cell marker *NANOG* was not detected and POU5F1 (Oct-4) levels were negligible. These results suggest a mixed population of cells enriched in ventral spinal neurons with the presence of some glial and progenitor cells, consistent with previous single-cell RNA-seq experiments.^45^ Importantly, the expression levels of cell type markers did not significantly differ between mutant and control samples, indicating a similar cell composition and differentiation propensity, as previously described.^43^

**Figure 1.**
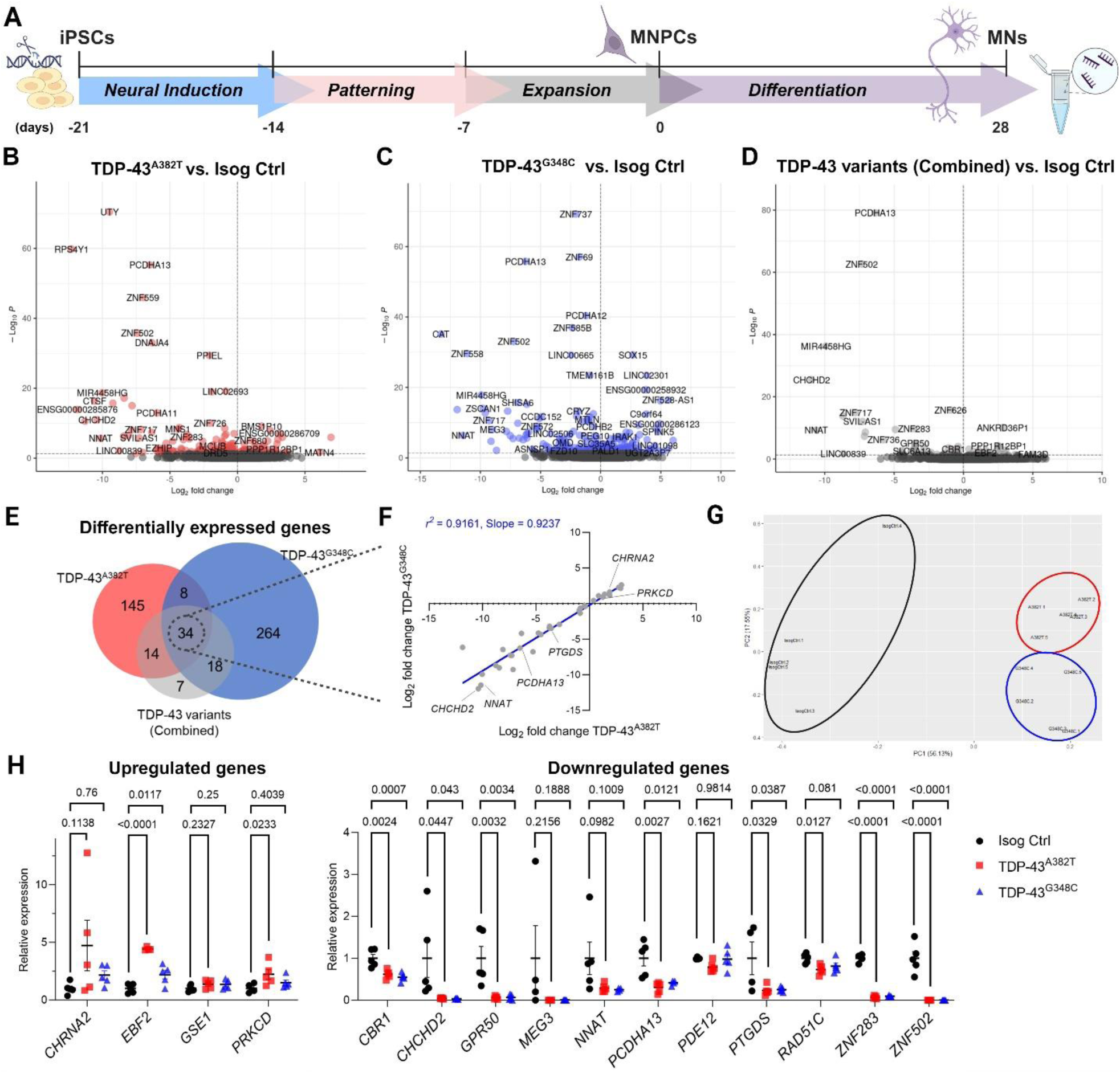
Differential gene expression analysis in TDP-43^A382T^ and TDP-43^G348C^ MNs. **(A)** Schematic representation of iPSCs differentiation into MN progenitor cells (MNPCs) and MNs for total RNA extraction. **(B-E)** Volcano plots **(B-D)** and Venn diagram **(E)** comparing differentially expressed genes (DEGs) in TDP-43 MNs differentiated for 28 days (4 weeks) relative to isogenic control. n=5 independent differentiations. **(F)** Scatter plot showing a strong correlation between fold changes of overlapping DEGs. **(G)** Principal component analysis (PCA) of the normalized gene expression of DEGs (combined contrast). **(H)** Validation of selected common DEGs by qPCR. Data shown as mean ± SEM. One-way ANOVA with Dunnett’s post hoc test. *P*-values shown. Significance was defined as *P*<0.05. n=5 independent differentiations.

### TDP-43^A382T^ and TDP-43^G348C^ MNs affect similar genes and pathways in human MNs

To identify transcriptional changes induced by *TARDBP* mutations, we performed differential gene expression analysis using DESeq2 (false discovery rate [FDR]<0.05) by contrasting each mutation to the isogenic control (**Figure 1, B and C**; **Supplemental Table S3**). We identified 201 differentially expressed genes (DEGs) in TDP-43^A382T^ MNs compared with control (137 downregulated genes and 64 upregulated genes), and 324 DEGs in TDP-43^G348C^ MNs compared with control (161 downregulated genes and 163 upregulated genes) (**Supplemental Figure 1, C and D**). The expression levels of the ALS-linked genes *SOD1*, *C9ORF72*, *FUS*, and *TARDBP* did not differ between TDP-43 and isogenic control samples (**Supplemental Figure 1B**). When comparing the genes dysregulated in TDP-43^A382T^ and TDP-43^G348C^ MNs, there was a significant overlap (*P*<0.05, hypergeometric test), revealing a total of 42 DEGs in common (**Supplemental Figure 1D**; **Supplemental Table S4**). Additionally, we combined the datasets from the two mutations and performed differential expression analysis against the isogenic control (TDP-43^A382T^+TDP-43^G348C^ vs. Isog Ctrl) (**Figure 1D**; **Supplemental Table S3**), with the goal of identifying DEGs with a shared direction of effect in TDP-43 samples. This analysis yielded a total of 73 DEGs. Among those, 14 and 18 genes were uniquely dysregulated in TDP-43^A382T^ and TDP-43^G348C^ MNs respectively in the prior analyses, while 34 genes were shared between the individual and combined contrasts (**Figure 1E**). There was a strong correlation between fold changes in shared differentially expressed genes (*r*^2^=0.9161, slope=0.9237, *P*<0.0001) (**Figure 1F**), indicating a similar magnitude of effect on gene expression by the two mutations. The convergence between TDP-43^A382T^ and TDP-43^G348C^ MNs was corroborated by findings from a principal-component analysis of DEGs (**Figure 1G**), where PC2 distinguished TDP-43^A382T^ and TDP-43^G348C^ samples into two distinct but closely related clusters.

We selected a subset of genes for validation by qPCR. We focused on genes showing an overlap between TDP-43^A382T^ and TDP-43^G348C^ MNs, and those related to cellular processes relevant to ALS and/or associated with ALS or more broadly to neurological disorders (**Supplemental Table S4**). We also took into consideration the expression levels (i.e., normalized counts) and the log fold changes between conditions. These strategies facilitated the selection of a total of 15 genes among those dysregulated in TDP-43 MNs. Differential expression of 10 genes was confirmed in either or both TDP-43 MN cultures in five additional RNA samples from a separate set of extractions (*EBF2*, *PRKCD*, *CRB1*, *CHCHD2*, *GPR50*, *PCDHA13*, *PTGDS*, *RAD51C*, *ZNF283*, *ZNF502*) (**Figure 1H**).

In addition to a shared gene signature, Gene Ontology (GO) enrichment analyses revealed that both mutants showed a dysregulation in genes with molecular functions related to ion binding, DNA binding, and transcription factor activity (**Figure 2A**), consistent with altered DNA/RNA-binding protein function. Additionally, genes dysregulated in TDP-43^G348C^ MNs showed a significant enrichment in the GO cellular component terms “Integral component of plasma membrane”, “intrinsic component of plasma membrane” and “synaptic membrane” as well as in GO biological processes related to cell adhesion (**Figure 2, B and C**). Interestingly, the “cell adhesion” category has similarly been reported to be enriched in several transcriptomic studies of ALS patient samples and iPSC-derived MN models.^22,46,47^

**Figure 2.**
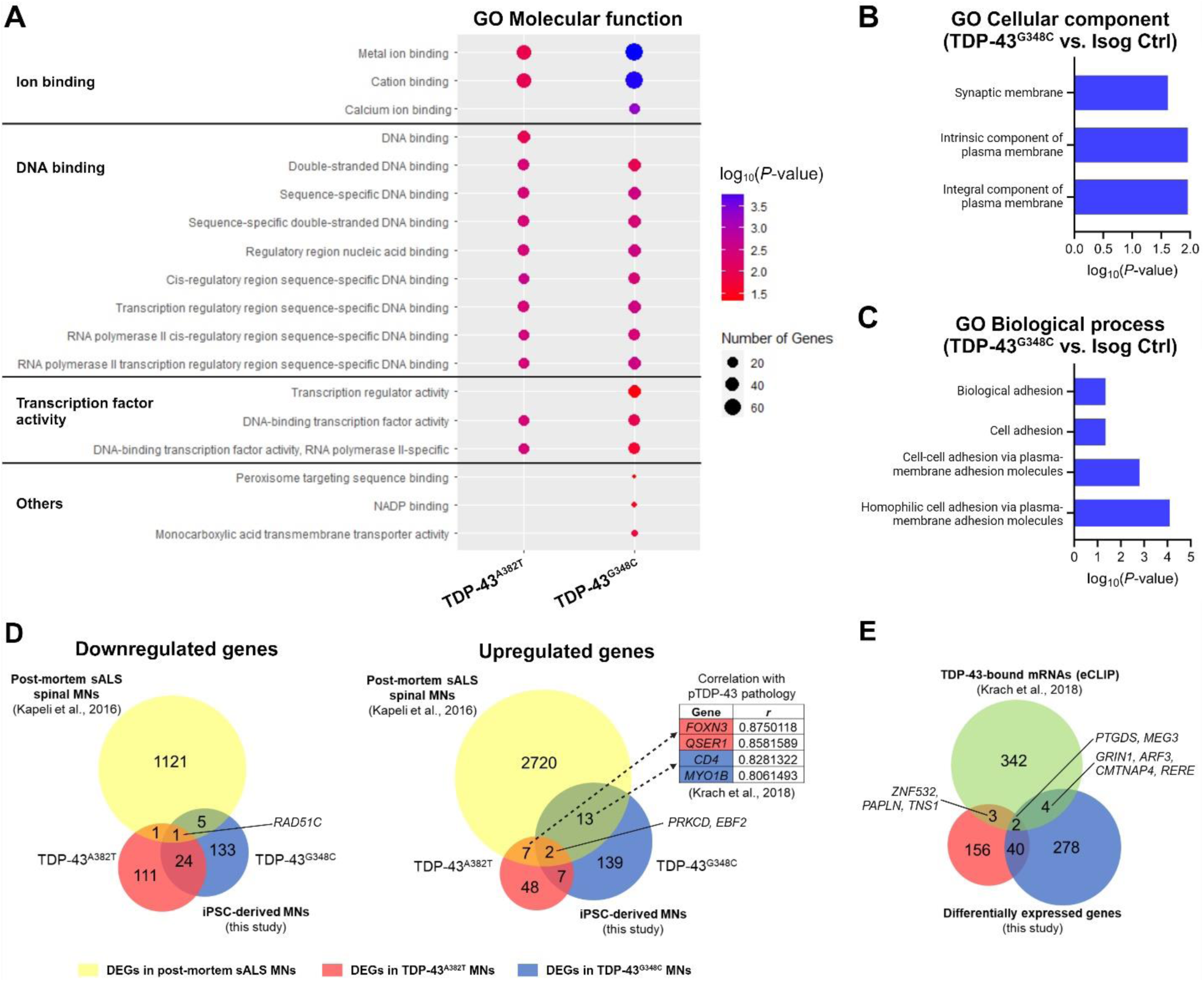
Functional characterization of differentially expressed genes. **(A)** Representative gene ontology (GO) terms of the “molecular function” category enriched in DEGs of TDP-43^A382T^ and TDP-43^G348C^ MNs. **(B** and **C)** Representative GO terms of the “cellular component” **(B)** and “biological process” **(C)** categories enriched in DEGs of TDP-43^G348C^ MNs. **(D)** Venn diagram comparing common DEGs between TDP-43 MNs and post-mortem sALS MNs. **(E)** Venn diagram comparing DEGs and a list of known TDP-43 mRNAs binding targets.

### TDP-43 MNs share gene expression changes with post-mortem sporadic ALS spinal MNs

To determine whether the genes affected by mutations in *TARDBP* resemble those dysregulated in sporadic ALS (sALS), we compared our RNA-seq data with a previously published dataset obtained from laser-captured spinal MNs from 13 sALS patients with TDP-43 pathology and 9 neurologically healthy individuals (**Figure 2D**).^48^ This study had identified a total of 3878 DEGs in sALS samples compared with controls (FDR<0.05); specifically, 2742 upregulated genes (70.1%) and 1128 downregulated genes (29.1%). With this gene set and ours, we tested the hypothesis that similar genes are affected in MNs from sALS patients and in iPSC-derived MNs with mutations in *TARDBP*. Two downregulated genes (*AMPH*, *RAD51C*) and nine upregulated genes (*EBF2*, *PRKCD*, *MME*, *FOXN3*, *ZNF532*, *HIF3A*, *QSER1, ZFHX3*) were common between sALS and TDP-43^A382T^ MNs. Six downregulated genes (*RAD51C*, *BMP3*, *PANK1*, *PEG10, MCTP1*) and 15 upregulated genes (*EBF2*, *PRKCD*, *PCGF3*, *RPS29*, *ZFAS1*, *PCDHGB7*, *PALD1*, *CD4*, *MYO1B*, *SOX13*, *LRRC34*, *SLCO1A2*, *PADI2*, *RGS9*, *CSRP2*) were shared between sALS and TDP-43^G348C^ MNs. Finally, three genes were shared between both mutants and sALS samples (*RAD51C*, downregulated and *PRKC2* and *EBF2*, upregulated). Interestingly, the expression of four shared upregulated genes (*FOXN3*, *QSER1*, *CD4, MYO1B*) was previously found to correlate with pTDP-43 pathology burden (*r*>0.8) in patient specimens.^20^ In summary, *TARDBP* mutant iPSC-derived MNs recapitulate some of the transcriptional dysregulation observed in sALS post-mortem samples.

### Few dysregulated genes are known TDP-43 targets

To further investigate the mechanisms underlying RNA changes induced by *TARDBP* mutations, we next considered a relationship between differential gene expression and mRNA binding by TDP-43. We turned to a previous study characterizing TDP-43 target mRNAs by enhanced crosslinking and immunoprecipitation (eCLIP) followed by RNA-seq in motor cortex specimens from sALS patients and neurologically healthy individuals.^20^ We compared our DEGs with a list of genes whose mRNAs were previously reported to be bound by TDP-43 (**Figure 2E**). We didn’t find a significant enrichment for TDP-43 targets within our datasets (TDP-43^A382T^ *P*<0.086; TDP-43^G348C^ *P*<0.177, hypergeometric test). Only a small fraction of DEGs were known TDP-43 targets (∼2,5% in TDP-43^A382T^ and ∼1.8% in TDP-43^G348C^). Two TDP-43-bound genes were dysregulated in both TDP-43 MN cultures (*PTGDS*, *MEG3*), while three and four TDP-43-bound genes were only dysregulated in TDP-43^A382T^ (*ZNF532*, *PAPLN*, *TNS1*) or TDP-43^G348C^ MNs (*GRIN1*, *ARF3*, *CNTNAP4*, *RERE*), respectively. Among dysregulated TDP-43-bound genes, the majority (6/9) were downregulated. Given that, to our knowledge, similar cross-linking experiments characterizing TDP-43-bound transcripts have not yet been performed in human spinal cord specimens or spinal MN cultures, additional RNA-protein interaction studies may be needed to reveal cell type-specific TDP-43 binding targets. In line with this idea, recent studies have shown that different cell environments modulate the RNA processing functions of TDP-^43.49,50^

### MicroRNA biogenesis is altered in mutant MNs

Given the roles of TDP-43 in miRNA biogenesis,^26,51^ we next interrogated whether *TARDBP* mutations could indirectly affect gene expression through changes in miRNA abundance. To this end, we performed miRNA profiling by small RNA sequencing to characterize miRNA abundance in TDP-43^A382T^ and TDP-43^G348C^ MN cultures. We conducted differential expression analyses (FDR<0.05) using the same contrast design used in the RNA-seq experiments (**Figure 3, A-C**; **Supplemental Table S5**). Differential expression analyses revealed a total of 40 and 47 miRNAs with an altered abundance in TDP-43^A382T^ and TDP-43^G348C^ MNs, respectively. The two mutants had largely overlapping dysregulated miRNAs (*P*<0.05, hypergeometric test), with 36 miRNAs in common (**Supplemental Figure S2A**; **Supplemental Table S6**). The combined analysis (TDP-43^A382T^+TDP-43^G348C^ vs. Isog Ctrl) identified a total of 46 dysregulated miRNAs, including the 36 miRNAs shared with the individual analyses (**Figure 3D**). Most miRNAs showed decreased abundance with a strong correlation in fold changes between the two mutants (*r*^2^=0.9201, slope=0.8254, *P*<0.0001) (**Figure 3E**), which could indicate altered miRNA biogenesis or excess degradation. Downregulation was confirmed in 1 out of 7 miRNAs selected for validation (has-miR-381-3p), although most miRNAs showed a clear trend toward decreased abundance relative to the isogenic control (**Figure 3F**).

**Figure 3.**
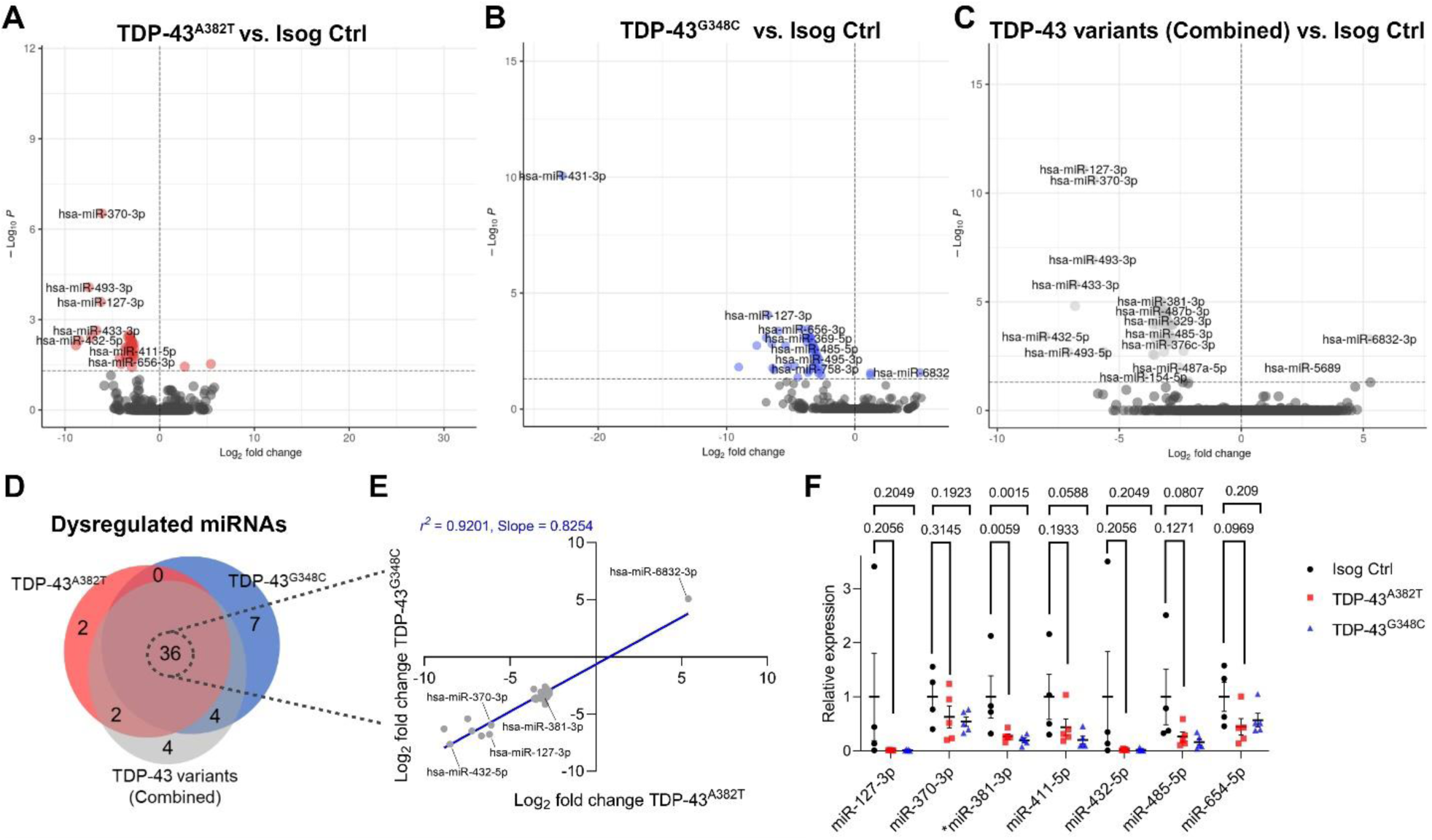
Differential miRNA expression analysis in mutant MNs. **(A-d)** Volcano plots **(A-C)** and Venn diagram **(D)** comparing differentially expressed miRNAs in TDP-43 MNs relative to isogenic control. n=5 independent differentiations. **(E)** Scatter plot showing a strong correlation between fold changes of overlapping dysregulated miRNAs. **(F)** Validation of selected common dysregulated miRNAs by qPCR. Data shown as mean ± SEM. One-way ANOVA with Dunnett’s post hoc test. *P*-values shown. Significance was defined as *P*<0.05. n=5 independent differentiations.

Given that TDP-43 can directly associate with pri-miRNA and pre-miRNA precursors during miRNA maturation,^26,51^ we queried the RBPmap database^52^ to search for TDP-43 binding motifs in the transcribed sequences of dysregulated miRNAs. We found one or more predicted TDP-43 binding sites in nearly half of the shared dysregulated miRNAs (17/36 miRNAs, **Supplemental Table S7**). Interestingly, we also noticed that the genomic coordinates of all 35 downregulated miRNAs mapped to the 14q32 miRNA cluster (**Supplemental Table S6**), which strongly implies that their expression is co-regulated. Accordingly, most of them were downregulated by the same magnitude (∼3 log fold-changes) in TDP-43 samples relative to the isogenic control in differential expression analyses (**Figure 3E**; **Supplemental Table S6**). Taken together, these results indicate that mutations in *TARDBP* lead to a decreased abundance of 14q32-encoded miRNAs.

### Functional consequences of dysregulated miRNA in mutant MNs

To explore the potential functional consequences of miRNA perturbances, we used miRgate^53^ to predict targets of shared dysregulated miRNAs (**Figure 4A**; **Supplemental Table S8**), filtering for target genes predicted by at least two out of five computational algorithms used by this database. Given that miRNAs can regulate gene expression post-transcriptionally through mRNA degradation, we considered an inverse correlation between dysregulated miRNAs and the expression levels of their predicted target mRNAs. The list of predicted genes targeted by downregulated miRNAs was compared to genes upregulated in mutant MNs determined by RNA-seq data (and vice versa). These mRNA/miRNA cross-analyses yielded a total of 13 and 30 genes identified as inversely correlated to miRNA abundance in TDP-43^A382T^ and TDP-43^G348C^ MNs, respectively, while a total of 6 genes were identified in the combined analysis (TDP-43^A382T^+TDP-43^G348C^ vs. Isog Ctrl) (**Figure 4B and Supplemental Figure S2B**). These results indicate that the detected changes in gene expression could be partially explained by impaired miRNA biogenesis. Interestingly, we noted that several negatively correlated genes encode protocadherins (**Supplemental Figure S2B**), which are cell-cell adhesion proteins enriched in the central nervous system (CNS).^54^ Upon re-examination of the RNA-seq data, we also found that DEGs were statistically enriched for the clustered protocadherin locus on chromosome 5 in both mutants (FDR<0.05) (**Supplemental Figure S2D**).

**Figure 4.**
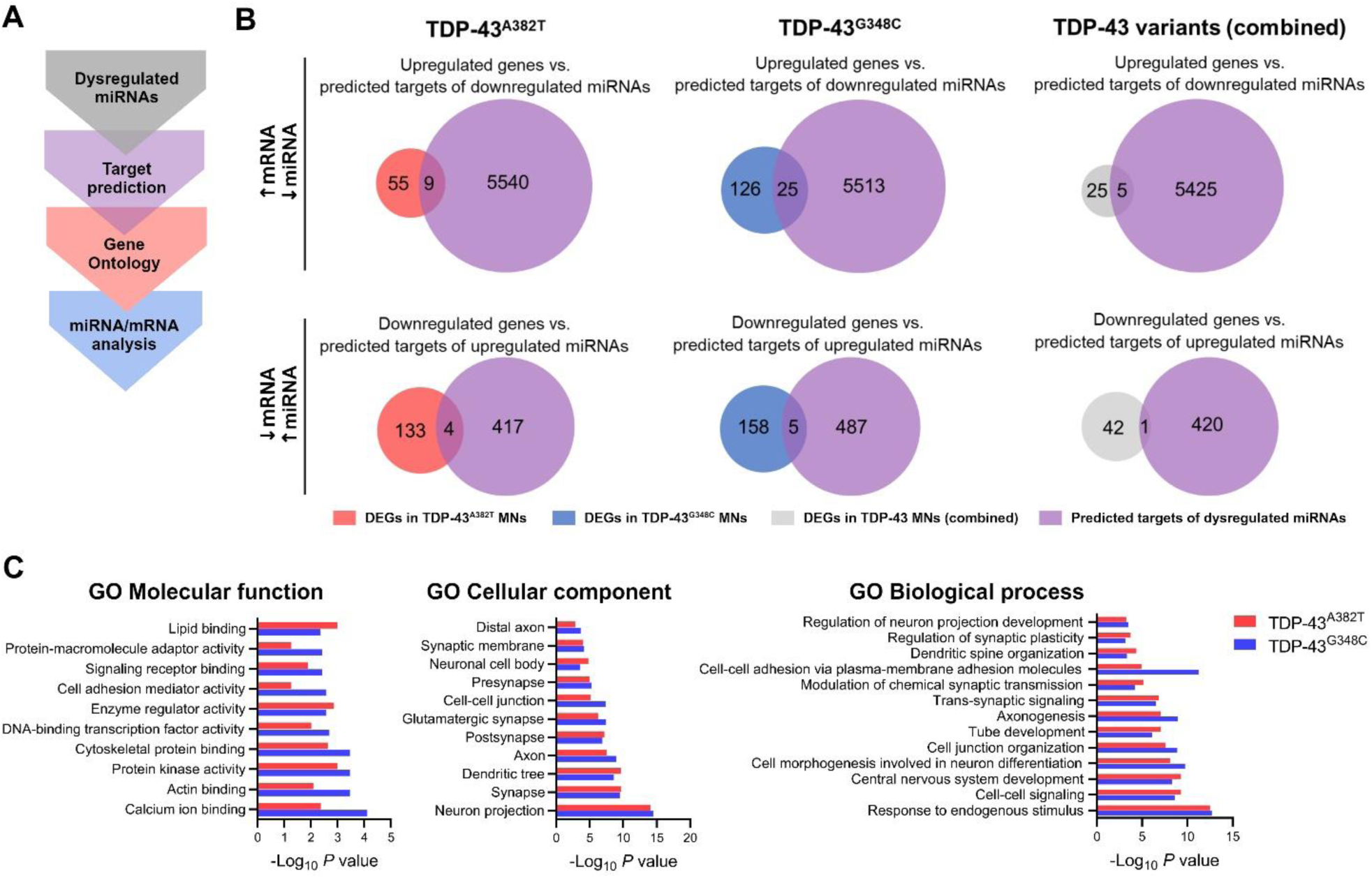
Functional characterization of dysregulated miRNAs. **(A)** Bioinformatics analysis workflow for functional characterization of dysregulated miRNAs. **(B)** Integrated mRNA/miRNA analysis comparing DEGs and predicted mRNA targets of dysregulated miRNAs in TDP-43 MNs. **(C)** Representative common enriched GO terms of predicted mRNA targets of dysregulated miRNAs.

Aside from targeted mRNA degradation, miRNAs can achieve RNA silencing through translational repression, with little to no influence on mRNA target abundance.^55^ To gain further insights into the pathways regulated by altered miRNA, we performed GO enrichment analyses of predicted target genes and investigated shared enriched GO terms per category. Overall, there was a common enrichment in GO terms related to synaptic function, cell adhesion, and neuronal development (**Figure 4C**). Furthermore, KEGG pathway analysis identified a common enrichment in pathways related to cell signaling, axon guidance, and neurodegeneration (**Supplemental Figure S2C**).

### *In silico* screen identifies compounds predicted to restore mutation-induced gene expression signatures

Next, we reasoned that mutation-induced gene expression profiles might represent a potential therapeutic opportunity. We posited that transcriptomic alterations may be reflective of underlying disease mechanisms and that normalizing gene expression may have beneficial effects. Thus, we utilized the CMap database^42^ (which comprises of transcriptional profiles induced from thousands of small molecules) to screen for compounds predicted to normalize gene expression changes caused by *TARDBP* mutations toward wild-type levels (**Figure 5A**). We used as inputs our lists of DEGs identified by RNA-seq and obtained as outputs lists of compounds accompanied by their τ scores, indicating the similarity or dissimilarity between the transcriptional changes they induce and the queried DEGs list (**Supplemental Tables S9 and S10**). The scores obtained from each queried DEG list were plotted to allow visualization of hits that are common between individual and combined datasets (**Figure 5B**). Among 16 top-compounds (score<-85), we ultimately selected for further investigation 6 candidate compounds from the lower left corner (i.e., predicted to reverse gene expression changes) with high scores in the combined dataset or at least one of the individual datasets (**Figure 5C**). To make this selection, we took into consideration the compound’s current approved therapeutic uses or investigational uses, known mechanisms of action, and blood-brain barrier penetrance.

**Figure 5.**
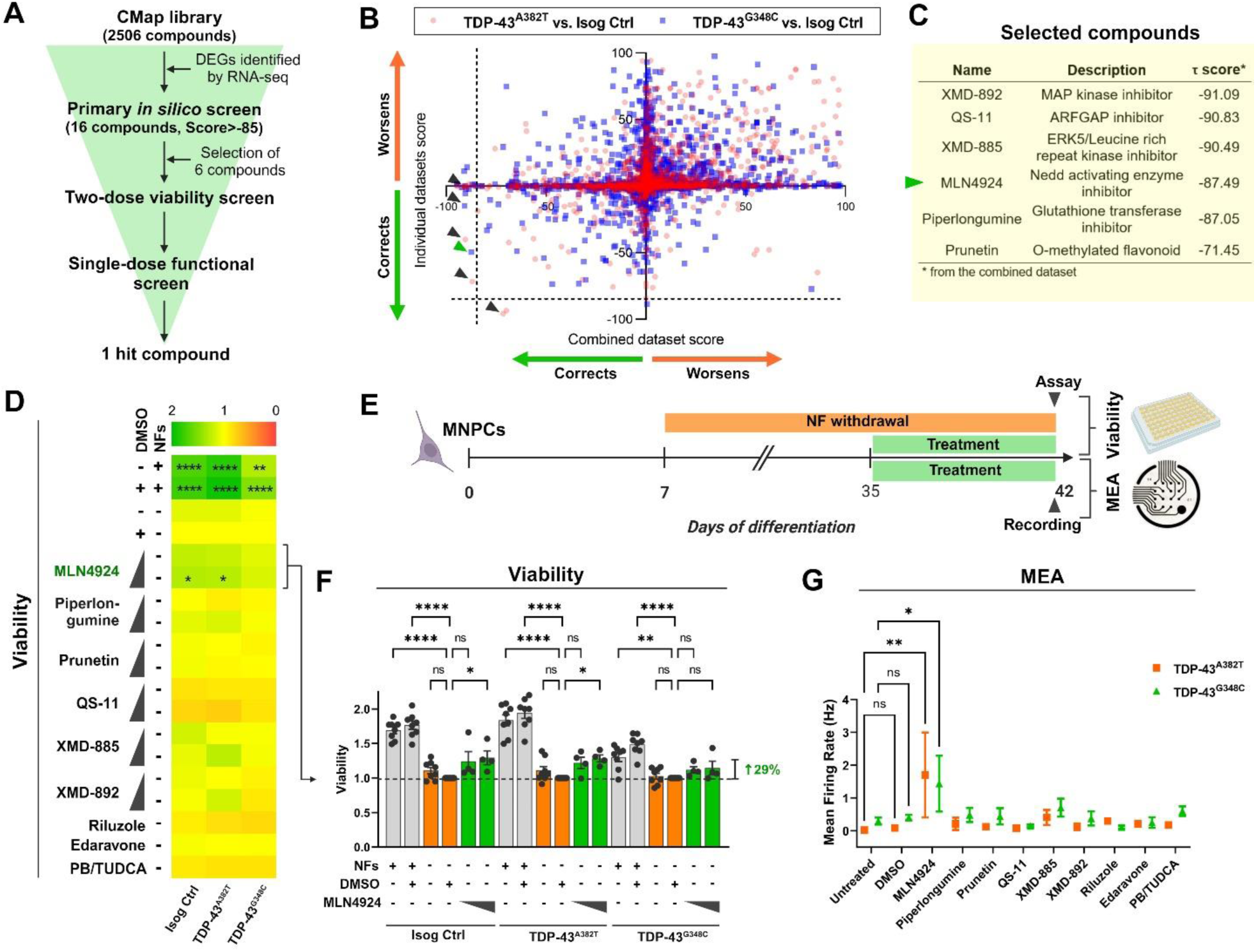
Transcriptome-based *in silico* and *in vitro* phenotypic screens identify one compound that ameliorates MN survival and activity. (A) Screening funnel used to identify candidate compounds. **(B)** Scatter plot of the τ scores from the individual (y-axis) versus combined (x-axis) datasets gene signatures. Each data point represents one compound of the CMap library. Arrow heads show selected compounds. **(C)** Description and τ scores of selected compounds. **(D)** Heatmap of mean MN survival relative to DMSO control after treatment with compounds (0.1 μM and 1.0 μM) in culture conditions without neurotrophic factors (NFs) supplementation. n=4 independent differentiations. One-way ANOVA with Sidak’s post hoc test. **(E)** Overview of *in vitro* phenotypic screens. **(F)** Bar graph of MN viability after treatment with MLN4924. n=4 independent differentiations. One-way ANOVA with Sidak’s post hoc test. **(G)** Mean firing rate of MNs treated with candidate compounds (1.0 μM) calculated from per-well mean values from n=4 independent differentiations. Two-way ANOVA with Dunnett’s post hoc test. Data shown as mean ± SEM. *p<0.05, **p<0.01., ***p<0.001, ****p<0.0001.

### Treatment with hit compound MLN4924 ameliorates MN viability and firing activity

In our next steps, we aimed to determine whether the six selected hit compounds could improve ALS-relevant phenotypic readouts, such as MN viability (**Figure 5A**). In previous work, we showed that removal of neurotrophic factors (NFs) supplementation from the differentiation medium significantly decreased the viability of MNs regardless of their genotype.^43^ Thus, we treated MNs cultured without NFs with two concentrations of the candidate compounds (0.1 μM and 1.0 μM) for 6 days to screen for compounds with neuroprotective properties (**Figure 5E**). We also added to our panel of compounds three FDA-approved drugs for the treatment of ALS, namely riluzole, edaravone, and the combination of sodium phenylbutyrate (PB) and tauroursodeoxycholic acid (TUDCA) (i.e., AMX0035). Following treatment, one compound (MLN4924) effectively improved viability of isogenic control and TDP-43^A382T^ MNs compared with the vehicle-treated condition at the highest tested concentration, with a similar trend observed in TDP-43^G348C^ MNs (Isog Ctrl: +29.3%, *P*=0.0352; TDP-43^A382T^: +28.4%, *P*=0.0472; TDP-43^G348C^: +14.5%, *P*=0.8681 by one-way ANOVA with Sidak’s multiple comparisons test) (**Figure 5, D and F**).

We previously showed using multi-electrode array (MEA) that TDP-43 MNs displayed decreased spontaneous neuronal activity compared with isogenic control MNs after several weeks in culture.^43^ In addition to viability, we thought it was important to also test for amelioration of this mutation-induced phenotype in response to treatment. Thus, we examined the effects of the six selected hit compounds on neuronal activity in an additional single-dose screen at the highest concentration (1.0 μM) (**Figure 5A**). For this assay, MNs were cultured in complete final differentiation medium (with NFs supplementation) to maintain optimal MN viability for electrophysiological recordings (**Figure 5E**). We found that MLN4924 treatment significantly increased the mean firing rate of TDP-43^A382T^ and TDP-43^G348C^ MN cultures compared to untreated cells (TDP-43^A382T^: +1.7 Hz, *P*=0.0091; TDP-43^G348C^: +1.5 Hz, *P*=0.0485 by two-way ANOVA with Dunnett’s multiple comparison test) (**Figure 5G**). Overall, we demonstrate that combining *in silico* and multi-phenotypic screens approaches enabled the identification of one compound able to improve neuronal viability and activity.

## Discussion

Accumulating evidence implicates RNA dyshomeostasis in the pathobiology of ALS. Here, we performed whole-transcriptome profiling of MNs differentiated from iPSCs with mutations in *TARBDP* coding for TDP-43^A382T^ or TDP-43^G348C^. Few studies previously conducted high-throughput transcriptomic analyses of *TARDBP* mutant iPSC models.^50,56,57^ Our study is the first to provide both mRNA and miRNA expression profiles with integrated mRNA/miRNA analyses, with the aim of increasing our understanding of TDP-43 pathobiology in ALS and identifying molecules able to restore disease-relevant phenotypes.

Our results highlight significantly overlapping RNA alterations in TDP-43^A382T^ and TDP-43^G348C^ MNs, indicating common pathogenic effects of these mutations on RNA processing. Importantly, our data points to several genes previously linked to ALS and other neurodegenerative disorders including frontotemporal dementia (FTD), Parkinson’s disease (PD), and Lewy body dementia (LBD). Among shared DEGs, the long non-coding RNA *MEG3*, a known TDP-43 target,^20,58^ was downregulated in both mutant datasets, and loss of TDP-43 binding to *MEG3* was previously observed in brains of subjects with FTD.^59^ The PD- and LBD-associated gene *CHCHD2*^60–62^ was also significantly downregulated in both mutants. Coiled-coil-helix-coiled-coil-helix domain containing 2 (encoded by *CHCHD2*) forms a complex with CHCHD10, another nucleus-encoded mitochondrial protein of the CHCHD-containing protein family, known to be genetically linked to ALS and FTD.^63–65^ Interestingly, *CHCHD10* was also significantly downregulated in TDP-43^A382T^ MNs. Previous studies have similarly reported loss of *CHCHD2* expression in TDP-43^M337V^ and TDP-43^Q331K^

iPSC-derived MNs^56^ and decreased *CHCHD10* expression in sALS patient-derived MNs.^66^ Additionally, *CHCHD2* and two other shared DEGs identified here (*PRKCD*, *MYH14*) were predicted to be candidate ALS genes in a recent machine learning study.^67^

Further comparison of our datasets with that of post-mortem sALS MNs revealed common transcriptomic alterations (e.g., *PRKCD)*,^48^ some previously correlated with TDP-43 pathology (*FOXN3*, *QSER1*, *CD4, MYO1B*),^20^ although the overlap was quite modest. We speculate that TDP-43 MNs may more closely resemble sALS MNs derived from iPSCs rather than their post-mortem counterparts. Indeed, residual surviving MNs recovered from post-mortem specimens are representative of the end-stage disease while iPSC-derived MNs, although less mature,^68^ might provide insights into the RNA changes occurring earlier in the disease course.

It is worth noting that TDP-43^A382T^ and TDP-43^G348C^ MNs did not exhibit a transcriptional signature of TDP-43 loss-of-function (implicating *STMN2*, *UNC13A*, *ELAVL3*, *PFKP*, *RCAN1*, *SELPLG*) as seen in knockdown studies and other ALS iPSC models,^17,18,50,69–72^ perhaps consistent with the preserved nuclear localization of TDP-43 in our cells.^43^ Thus, it appears that the TDP-43^A382T^ and TDP-43^G348C^ variants impact RNA processing by distinct mechanisms.

We explored the possibility that the *TARDBP* mutations studied could impact gene expression through altered miRNA biogenesis. miRNAs are highly expressed in the CNS and have been shown to be dysregulated in ALS patient specimens.^73–79^ We found a marked overlap in dysregulated miRNAs in both mutant MNs with concordant fold changes and directionality, some of which were previously linked to ALS. Specifically, miR-329-3p was dysregulated in muscle tissue of SOD1^G93A^ mice,^78^ while dysregulation of miR-370-3p, miR-409-3p, and miR-495-3p was also documented in iPSC-derived MNs with homozygous FUS^P517L^ mutations.^80^ Other dysregulated miRNAs, namely miR-432-5p, miR-323a-3p and miR-453, were predicted here to target *ALS2*, associated with juvenile ALS.^81,82^ Additionally, the ALS-associated gene *SARM1*^83,84^ was predicted to be regulated by miR-432-5p, mir-370-3p, miR-453, miR-495-3p, and miR-654-5p, all miRNAs that were downregulated in our study. Thus, our work supports the view that miRNAs are involved in ALS pathophysiology.

Strikingly, most altered miRNAs mapped to the 14q32 miRNA cluster, implying that their expression is regulated by a common transcriptional or epigenetic process. Members of miRNAs clusters typically share similar biological functions by targeting the same sets of genes or functionally related genes of the same pathways.^85^ The miRNAs of the 14q32 cluster (also known as the miR379–410 or C14MC cluster) are highly conserved, brain-enriched miRNAs which are involved in many aspects of neuronal development and function including neuronal migration, neurite formation, and synaptic plasticity.^86,87^ Accordingly, we found several differentially regulated miRNA/mRNA pairs implicating protocadherins, which play key roles in neural circuit formation.^54^ The miRNAs of this cluster were also previously associated with brain disorders including epilepsy,^88,89^ schizophrenia,^90,91^ autism spectrum disorder,^92,93^ and brain cancers.^94,95^ Interestingly, miRNAs of this cluster were shown to be selectively exported via exosomes,^96^ and several have been detected in the serum,^97^ suggesting they could constitute promising biomarkers. In fact, levels of the 14q32-encoded miRNA miR-127-3p were altered in cells of this study as well as in the serum of sporadic and familial ALS patients in two previous studies.^98,99^

In addition to biomarkers, miRNAs are also increasingly regarded as potential therapeutic targets, given that miRNAs can modulate hundreds of targets simultaneously.^100^ In future work, individual miRNA mimics or synthetic miRNA clusters^101^ may be employed to modulate levels of dysregulated miRNAs and their targets to assess phenotypic rescue. Such therapeutic strategies capable of acting more broadly on multiple cellular pathways, rather than on a few targets, may be more likely to have a clinically significant impact on the disease course. However, an important consideration for miRNA replacement therapy is the potential for undesirable off-target effects, which can be hard to predict or avoid^102^ as well as the invasive nature of the treatments (i.e., intrathecal injections) that are not without risk for patients.

Small orally bioavailable molecules able to improve the function and viability of MNs may be promising ALS therapies, given their ease of administration. Thus, we leveraged the transcriptomic profiles of TDP-43 MNs to search for compounds predicted to correct gene expression using the CMap database. This drug discovery strategy was previously applied to several human conditions including aortic valve disease,^103,104^ non-alcoholic steatohepatitis,^105^ epilepsy,^106–108^ schizophrenia,^109^ neurodevelopmental disorders,^41^ and FTD,^110^ yet remains largely unexplored in the ALS field.

Among top-scoring compounds, a number of compounds had pharmacological effects potentially relevant for the treatment of ALS. Indeed, some of them were known to act in the CNS for the treatment of neurological symptoms (e.g., neuroleptic/antipsychotic, anticonvulsant) and/or were ligands of neurotransmitter receptors (e.g., dopamine receptor, serotonin receptor), namely sulpiride, QS-11, GR-46611, and CS-110266. Sulpiride is a neuroleptic and all hit compounds of a previous large-scale chemical screen in TDP-43^A315T^ worms and TDP-43^G348C^ zebrafish were neuroleptics.^111^ Additionally, while there is evidence of increased MAP kinase signaling in ALS patients as well as in animal and iPSC models,^57,112–115^ two compounds identified here were MAP kinase inhibitors (XMD-892, XMD-885). Finally, XMD-892^116^ and other compounds (tacedinaline,^117–119^ MLN4924,^120^ piperlongumine,^121,122^ prunetin^123^) have also been under investigation for their antioxidant, anti-inflammatory, anti-ischemic, neuroprotective, and/or antitumour properties.

*In silico* compound prediction was followed by two *in vitro* screens based on viability and hypoactivity phenotypes described in previous work.^43^ We tested a panel of 6 top-scoring compounds along with three approved ALS drugs: riluzole, edaravone, and PB/TUDCA. These efforts led to the identification of MLN4924, which effectively improved neuronal viability and firing activity. Perhaps surprisingly, treatments with the ALS drugs did not show similar neuroprotective effects at the concentrations used. MLN4924 is a NEDD8-activating enzyme inhibitor, which prevents the conjugation of the ubiquitin-like protein NEDD8 on protein substrates, a post-translational modification (PTM) known as NEDDylation. Given that PTMs can regulate protein-protein interactions and liquid-liquid phase separation behavior,^124^ we speculate that NEDDylation inhibition by MLN4924 may modulate the formation of RNP condensates in MNs, a process central to several RNA processing functions of TDP-43. In fact, modulating the NEDDylation pathway has been linked to the assembly and disassembly of stress granules.^125–127^ NEDDylation has also been shown to promote nuclear protein aggregation under proteotoxic stress conditions.^128^ These results and the apparent neuroprotective effects of MLN4924 in our iPSC models point to a potential modulating role of the NEDDylation pathway in TDP-43 biology.

In summary, we showed that mutations in *TARDBP* coding for TDP-43^A382T^ and TDP-43^G348C^ led to common alterations in the abundance of mRNAs and 14q32-encoded miRNAs in iPSC-derived MNs, with integrated mRNA/miRNA analysis pointing to defects in cell-cell adhesion and synaptic function. This study demonstrates the utility of whole-transcriptome profiling not only to shed light on perturbed pathways that may be relevant to disease, but also to predict potential therapeutic candidates using a transcriptome reversal paradigm. *In silico* prediction of MLN4924 translated into improvement of disease-relevant phenotypes *in vitro* in our models, demonstrating the potential of this strategy for drug discovery in ALS and other neurodegenerative diseases.

## Methods

### iPSC lines and culture

A summary of the iPSC lines used can be found in **Supplemental Table S1**. iPSCs were maintained on dishes coated with Matrigel (Corning Millipore; Cat#354277) in mTeSR1 (StemCell Technologies; Cat#85850) with daily media change and passaged at 80% confluence using Gentle Cell Dissociation Reagent (StemCell Technologies; Cat#07174). Cultures were routinely tested for mycoplasma using the MycoAlert Mycoplasma Detection kit (Lonza; Cat#LT07-318).

### Differentiation of MNs from iPSCs

IPSCs were first induced into MN progenitor cells (MNPCs) by dual-SMAD signaling inhibition and ventral neural patterning as previously described.^129^ Banked MNPCs were thawed onto dishes coated with 10 µg/ml Poly-L-ornithine (PLO) (Sigma-Aldrich; Cat#P3655) and 5 µg/ml laminin (Sigma-Aldrich; Cat#L2020) in “expansion medium” composed of basic neural medium (1:1 mixture of DMEM/F12 medium (Gibco; Cat#10565–018) and Neurobasal medium (Life Technologies; Cat#21103–049), 0.5X N2 (Life Technologies; Cat#17502–048), 0.5X B27 (Life Technologies; Cat#17504–044), 0.5X GlutaMAX (Gibco, Cat#35050-061), 1X antibiotic-antimycotic (Gibco, Cat#15240–062), and 100 µM ascorbic acid (Sigma-Aldrich; Cat#A5960)) supplemented with 3 µM CHIR99021 (Selleckchem; Cat#S2924), 2 µM SB431542 (Selleckchem; Cat#S1067), 2 µM DMH1 (Selleckchem; Cat#S7146), 0.1 µM retinoic acid (RA, Sigma-Aldrich; Cat#R2625), 0.5 µM purmorphamine (PMN, Sigma-Aldrich; Cat#SML-0868), 0.5 mM valproic acid (VPA, Sigma-Aldrich; Cat#P4543)) and 10 µM ROCK inhibitor Y-27632 (Selleckchem; Cat#S1049) (for the first 24 h), and were allowed to recover up to 6 days with medium fully changed every other day.

For final MN differentiation, MNPCs were dissociated as single cells with Accutase (StemCell Technologies; Cat#07922) and were seeded on dishes coated with 10 µg/ml PLO and 5 µg/ml laminin (Life Technologies; Cat#23017-015) in “final differentiation medium” composed of basic neural medium supplemented with 0.5 µM RA, 0.1 µM PMN, 0.1 µM Compound E (CpdE, StemCell Technologies; Cat#73954), and 10 ng/ml insulin-like growth factor-1 (IGF-1, Peprotech; Cat#100–11), brain-derived neurotrophic factor (BDNF, Peprotech; Cat#450–02) and ciliary neurotrophic factor (CNTF, Peprotech; Cat#450–13)), with half-changes at least every other week. Alternatively, for western blot, viability, and MEA experiments, MNPCs were first passaged at a 1:3 ratio in “priming medium” composed of basic neural medium supplemented with 0.5 µM RA and 0.1 µM PMN with medium changed every other day for 6 days before they were plated for final differentiation as described above.

### Library preparation and sequencing

Total RNA was extracted from MN cultures using the miRNeasy micro kit (Qiagen; Cat#217004) with DNase treatment (Qiagen; Cat#79256) following the manufacturer’s instructions. Samples with RNA integrity numbers above 6.5 determined by a Bioanalyzer (Agilent) were used for library preparation using the NEB mRNA stranded library preparation kit (for RNA-seq) and the NEB small RNA library preparation kit (for small RNA-seq). Libraries were prepared and sequenced at Genome Québec (Montréal, Canada) on a NovaSeq 600 sequencing system with paired-end 100 bp (PE100) strategy.

### RNA-seq analysis

Sequencing files were analyzed using the GenePipes RNA-seq pipeline 3.6.0.^130^ Briefly, raw reads were clipped for adapter sequence, trimmed for minimum quality (Q30) in 3’ and filtered for minimum length of 32 bp using Trimmomatic.^131^ Surviving read pairs were aligned to the human genome assembly hg38 by the ultrafast universal RNA-seq aligner STAR^132^ using the recommended two passes approach. Aligned RNA-seq reads were assembled into transcripts and their relative abundance was estimated using Cufflinks.^133^ Differential expression analyses were conducted using DESeq2^134^ with a false discovery rate of 5% with no cut-off on log fold-change. Volcano plots were created using the Molecular and Genomics Informatics Core (MaGIC) Volcano Plot Tool, available at https://volcano.bioinformagic.tools/.

### miRNA profiling

Sequencing datasets were analyzed with a modified version of the workflow proposed by Potla et al.^135^ We adapted this workflow to analyze reads without UMI’s, single end and paired end readsets, and multiple readsets per sample (this workflow is available at https://github.com/neurobioinfo/miRNA_workflow). After quality control with FastQC (http://www.bioinformatics.babraham.ac.uk/projects/fastqc/), we used cutadapt to remove reads shorter than 15 nt and larger than 32 nt. We used Bowtie v1.3^136^ to align readsets to the miRBase database v22^137–141^ and to the human reference genome v104. In order to maximize the number of alignments and to detect rare miRNAs, we used a seed length 20 (Bowtie option -l 20) and allowed 3 missmatches within the seed (Bowtie option -n 3). MiRNAs aligned to miRBase were counted with Samtools idxstats.^142^ MiRNAs aligned to the reference genome were filtered with TAGBAM from BEDTools^143^ and counted with a locally developed script that optimizes counting time (https://github.com/neurobioinfo/miRNA_workflow). Finally, the local workflow adds miRNAs aligned to miRBase to those aligned to the reference genome for each sample, and generates a miRNA counts matrix with all samples counts. Before differential expression analyses with the R package DESeq2,^134^ the count matrix was pre-filtered to keep miRNAs having at least 10 counts. DESeq2 estimates dispersions using the Cox-Reid method.^144^ It uses negative binomial GLM fitting for log2 fold changes estimation and the Wald statistics for hypothesis testing. We used a false discovery rate of 5% with no cut-off on log2 fold change.

### Gene ontology and KEGG enrichment analysis

Terms from the Gene Ontology were tested for enrichment with the ShinyGO online tool, available at http://bioinformatics.sdstate.edu/go/.

### miRNA target prediction

Prediction of miRNA targets was performed using miRGate,^53^ which uses five different public prediction algorithms. We considered target genes predicted by at least two algorithms.

### Quantitative PCR

Expression levels of genes of interest were quantified using qPCR. cDNA synthesis was performed with 500 ng of total RNA using the M-MLV Reverse Transcriptase kit (Thermo Fischer Scientific; Cat#28025013) in a total volume of 40 μL. To quantify expression levels of selected miRNAs, cDNA synthesis was performed using 10 ng of total RNA with the TaqMan® Advanced miRNA cDNA Synthesis Kit following the manufacturer’s instructions. Real-time qPCR reactions were set up in triplicates with the Applied Biosystems Applied Biosystems SYBR green (Applied Biosystems; Cat#A25778) or TaqMan® Fast Advanced Master Mix (Applied Biosystems; Cat#A44360) and run on a QuantStudio 5 Real-Time PCR system (Applied Biosystems Cat#A28140). Primers and TaqMan® probes references are provided in **Supplemental Table S2**. Levels of the endogenous control miRNA hsa-miR-191-5p^145^ and the geometric mean of the endogenous control genes *18S (RN18S1)*, *HPRT1*, *PPIA* and *POLR2A* were used for normalization. Normalized expression was displayed relative to the relevant control condition.

### *In silico* screen with the CMap database

The lists of down/upregulated genes of each contrast were used as inputs in the CMap database^42^ to assess the similarity (or dissimilarity) between the query gene set and gene expression profiles induced by a library of small molecules. The obtained output is a list of the small molecules rank-ordered by their CMap connectivity score (τ), a normalized metric ranging from -100 to 100. A negative score indicates opposing profiles between the small molecule and the query gene set, thereby predicting the small molecule to reverse gene expression changes. In contrast, a positive score indicates similar profiles.

### Two-dose viability screen

MNPCs were plated as 15,000 cells per well in opaque white optical 96-well plates coated with PLO/laminin. Cells were cultured in final differentiation medium for one week, after which they were treated with 1 μM cytosine arabinoside (AraC, Sigma-Aldrich; Cat#C6645) overnight (∼17 h) to eliminate any residual proliferating cells. The next day, medium was fully changed to final differentiation medium with or without supplementation of neurotrophic factors (NFs) (i.e., BDNF, CNTF, and IGF-1), with half-changes every other week until initiation of the assay. After 5 weeks of final differentiation, MNs were treated with the candidate compounds MLN4924, piperlongumine, prunetin, QS-11, XMD-885, XMD-892 at the final concentrations of 0.1 µM and 1.0 µM, or with 3 µM riluzole, 0.025 µM edaravone or 250 µM PB/10 µM TUDCA for 6 days with medium renewed every second day. After treatment, viability was assessed with an ATP-based chemiluminescent assay (Cell Titer-Glo®, Promega; Cat#G7570) following the manufacturer’s instructions. The luminescence readings were acquired using a GloMax Microplate Reader (Promega). The percentage of viability was determined by normalizing the raw luminescence values to those of the vehicle condition (DMSO-treated wells, medium without NFs) for each cell line.

### Single-dose functional screen

To assess the effect of compounds on neuronal activity, treatments were performed in MNs differentiated for 5 weeks in 24-well MEA plates (Axion Biosystems) in complete final differentiation medium. MNs were treated with candidate compounds for 6 days at a final concentration of 1.0 µM or with the ALS drugs as described above. After treatment, MEA recordings and analyses were carried out as previously described.^43^ Briefly, MNs were incubated in freshly prepared pre-warmed carbonated artificial cerebrospinal fluid (aCSF)^146^ for at least 1 hour. Spontaneous activity was recorded for 5 min in a Maetro Edge MEA system (Axion Biosystems) and the AxiS v.66465 software (Axion Biosystems).

### Statistics

Biological replicates were defined as independent differentiations. Hypergeometric testing was performed to compare the lists of genes and miRNAs using the following web app: http://nemates.org/MA/progs/overlap_stats.html. For qPCR experiments, Grubbs’ test was used to determine significant outliers. Statistical analyses were performed with the GraphPad Prism 9.3.0 software. Data distribution was assumed to be normal although this was not formally tested. Differences between mutant and control were analyzed using one-way or two-way analysis of variance (ANOVA) tests. Means and standard errors of the mean were used for data presentation. Significance was defined as *p*<0.05.

### Study approval

The use of human cells in this study was approved by McGill University Health Center Research Ethics Board (DURCAN_iPSC / 2019-5374).

### Data availability

The RNA-seq and small RNA-seq datasets generated as part of this study will be made accessible through GEO.

## Author contributions

Conceptualization, S.L., G.M., M.C., and T.M.D.; Methodology, S.L., G.M., L.G., M.C., and T.M.D.; Software, G.J.A. and D.S.; Validation, S.L., A.N.J.; Formal analysis, S.L., G.J.A. and D.S.; Investigation, S.L., A.S., A.N.J., M.J.C.-M., N.A., A.K.F.-F., G.H.; Resources, M.C., G.M., and T.M.D.; Data curation, S.L.; Writing— original draft, S.L.; Writing—review and editing, S.L., G.M., M.C., and T.M.D.; Visualization, S.L., G.M., M.C. and T.M.D.; Supervision, G.M., M.C. and T.M.D.; Project Administration, S.L., M.C., and T.M.D.; Funding acquisition, T.M.D. All authors have read and agreed to the published version of the manuscript.

## Acknowledgements

We acknowledge Dr. Rhalena A. Thomas for guidance on sample preparation for RNA-seq. We are also grateful to Dr. Gary A.B. Armstrong and Dr Vincent Soubannier for creative discussions and their advice.

S.L. was supported by the Faculty of Medicine and Health Sciences of McGill University. This work was supported by the Canada First Research Excellence Fund, awarded through the Healthy Brains, Healthy Lives initiative at McGill University; the CQDM FACs program; and the US Department of Defense ALS Research Program. All figures and schematics were created with BioRender.com.

