## Supplemental Material for "Transcriptome-based screening in TARDBP/TDP-43 knock-in motor neurons identifies the NEDD8-activating enzyme inhibitor MLN4924"

### Table of Contents


#### I. Supplemental Figures

Figure S1. Transcriptomic profiling of TDP-43$^{A382T}$ and TDP-43$^{G348C}$ MNs

Figure S2. miRNAs profiling and integrated miRNA/mRNA analysis highlight protocadherin-coding genes

#### II. Supplemental Tables

Table S1. Overview of iPSC lines used

Table S2. List of primers and TaqMan probes

Table S3. Differentially expressed genes from RNA-seq experiments *(Excel spreadsheet)*

Table S4. Shared differentially expressed genes

Table S5. Differentially expressed miRNAs from small RNA-seq experiments *(Excel spreadsheet)*

Table S6. Shared dysregulated miRNAs and genomic location

Table S7. Predicted TDP-43 binding sites by the RBPmap database

Table S8. Predicted targets of dysregulated miRNAs by the miRGate database *(Excel spreadsheet)*

Table S9. Top-scoring predicted compounds by the CMap database

Table S10. CMap τ scores outputs *(Excel spreadsheet)*

#### IV. Supplemental References

### I. Supplemental Figures

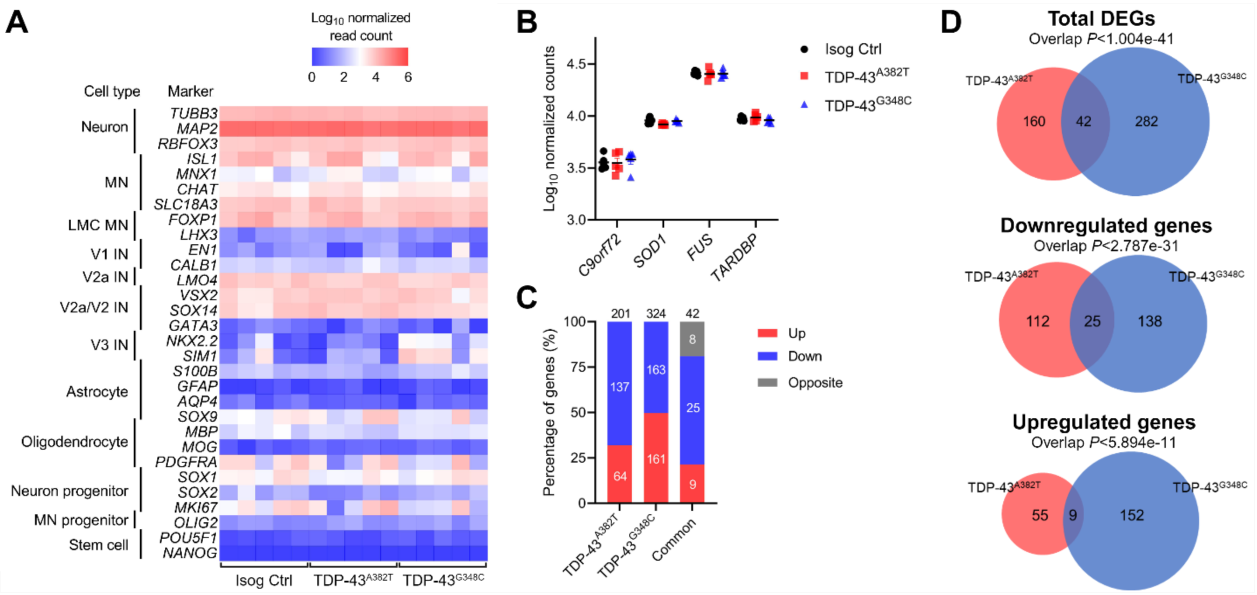


**Figure S1. Transcriptomic profiling of TDP-43$^{A382T}$ and TDP-43$^{G348C}$ MNs.**
**(A)** Heatmap of normalized counts of cell type markers determined by RNA-seq. **(B)** Normalized counts of ALS-associated genes. **(C and D)** Stacked bar graph **(C)** and Venn diagram **(D)** comparing differentially expressed genes (DEGs) in TDP-43 MNs relative to isogenic control. n=5.

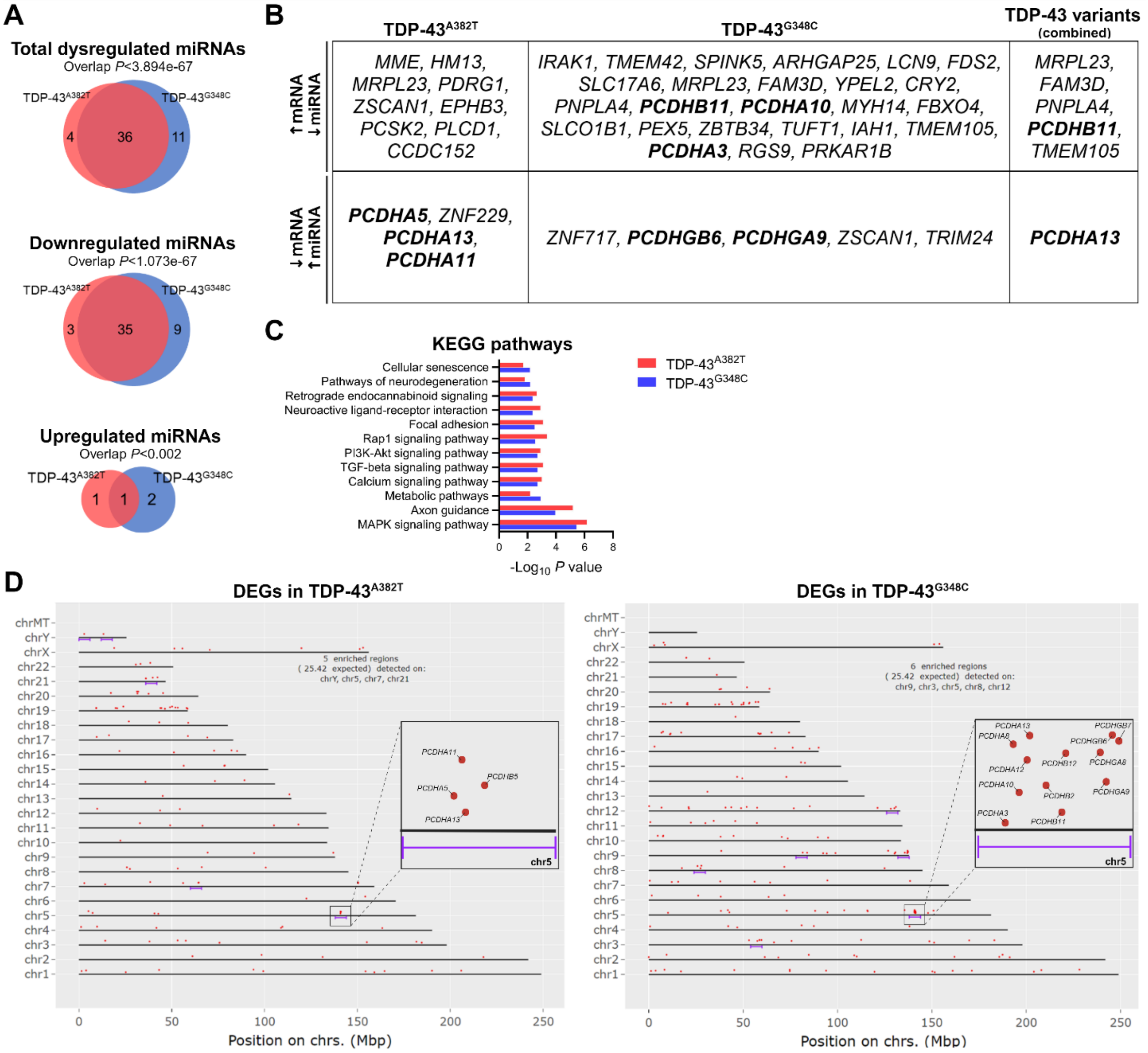


**Figure S2. miRNAs profiling and integrated miRNA/mRNA analysis highlight protocadherin-coding genes.**

**(A)** Venn diagram comparing differentially expressed miRNAs in TDP-43 MNs relative to isogenic control. n=5. **(B)** Differentially expressed genes (DEGs) predicted to be targeted by dysregulated miRNAs. **(C)** KEGG pathway analysis of predicted target genes of dysregulated miRNAs. **(D)** Genomic regions of statistical DEGs enrichment (FDR<0.05) determined by the ShinyGO web app. DEGs are enriched at the clustered protocadherin gene locus (chr5q31).

### II. Supplemental Tables

**Supplemental Tables S3, S5, S8, and S10** are provided as separate excel spreadsheets.

**Table S1. Overview of iPSC lines used**

| Cell line ID | ALS mutation | Sex | Age | Ethnicity | Primary cell line | Reprogramming method | Reference |
|---|---|---|---|---|---|---|---|
| AIW002-02 | None | Male | 37 | Caucasian | PBMC | Sendai virus | Chen et al. 2021[1] |
| *TARDBP* A382T/AIW002-02 | p.A382T | Male | 37 | Caucasian | Knock-in | n/a | Lépine et al. 2023[2] (preprint) |
| *TARDBP* G348C/AIW002-02 | p.G348C | Male | 37 | Caucasian | Knock-in | n/a | Lépine et al. 2023[2] (preprint) |

* PBMC, peripheral blood mononuclear cell; n/a, not applicable.

**Table S2. List of primers and TaqMan probes**

| Gene/miRNA | Reference |
|---|---|
| *18S (RN18S1)* | Hs.PT.39a.22214856.g |
| *HPRT1* | Hs.PT.58v.45621572 |
| *PPIA* | Hs.PT.58v.38887593.g |
| *POLR2A* | Hs.PT.39a.19639531 |
| *ZNF502* | Hs.PT.58.24576237 |
| *MEG3* | Hs.PT.58.25426100 |
| *PTGDS* | Hs.PT.58.2258847 |
| *ZNF283* | Hs.PT.58.27732356 |
| *RAD51C* | Hs.PT.58.40910176 |
| *PCDHA13* | Hs.PT.58.27085852.g |
| *GPR50* | Hs.PT.58.39777811 |
| *NNAT* | Hs.PT.58.24260117.g |
| *CHCHD2* | Hs.PT.58.38445645 |
| *PDE12* | Hs.PT.58.19439493 |
| *CBR1* | Hs.PT.56a.22365466 |
| *GSE1* | Hs.PT.58.40478286 |
| *EBF2* | Hs.PT.58.818465 |
| *PRKCD* | Hs.PT.58.21209513 |
| *CHRNA2* | Hs.PT.58.20688322 |
| hsa-miR-191-5p | 477952_mir |
| hsa-miR-432-5p | 478101_mir |
| hsa-miR-654-5p | 478368_mir |
| hsa-miR-381-3p | 477816_mir |
| hsa-miR-485-5p | 478126_mir |
| hsa-miR-411-5p | 478086_mir |
| hsa-miR-127-5p | 477889_mir |
| hsa-miR-370-3p | 478326_mir |

**Table S4. Shared differential expressed genes**

| Gene symbol** | Gene name | Functions/Comments*** | LogFC | |
|---|---|---|---|---|
| | | | p.A382T | p.G348C |
| ***Upregulated*** | | | | |
| *CYCSP49* | CYCS pseudogene 49 | Pseudogene | 2.925 | 2.149 |
| *PPP1R12BP1* | Protein phosphatase 1 regulatory subunit 12B pseudogene 1 | Pseudogene | 2.743 | 2.291 |
| *MRPL23* | Mitochondrial ribosomal protein L23 | Mitochondrial translation and metabolism of proteins | 1.417 | 1.324 |
| ***GSE1*** | Gse1 coiled-coil protein | May be subunit of a BRAF35-HDAC (BHC) histone deacetylase complex; overexpressed in breast cancer | 0.401 | 0.428 |
| *ANKRD36P1* | Ankyrin repeat domain 36 pseudogene 1 | Pseudogene | 2.990 | 2.644 |
| ***EBF2*** | EBF transcription factor 2 | Transcription factor | 1.897 | 1.186 |
| *NUTM2F* | NUT family member 2F | Protein-coding gene of unknown function | 0.852 | 1.084 |
| ***PRKCD*** | Protein kinase C delta | Related to autophagy and apoptosis; associated with various human diseases; predicted ALS gene[3] | 1.237 | 0.922 |
| ***CHRNA2*** | Cholinergic receptor nicotinic alpha 2 subunit | Nicotinic acetylcholine receptor subunit widely expressed in the brain; associated with epilepsy | 1.899 | 1.637 |
| ***Downregulated*** | | | | |
| *ZNF626* | Zinc finger protein 626 | Predicted to enable DNA-binding transcription factor activity | -0.949 | -0.925 |
| ***ZNF502*** | Zinc finger protein 502 | Predicted to enable DNA-binding transcription factor activity | -7.415 | -7.231 |
| *LINC00839* | Long intergenic non-protein coding RNA 839 | lncRNA | -8.769 | -8.667 |
| *ENSG00000285876* | Novel transcript | lncRNA | -11.862 | -6.14 |
| *ENSG00000269993* | Novel transcript | lncRNA | -3.728 | -3.179 |
| ***MEG3*** | Maternally expressed 3 | lncRNA associated with rare syndromes; known TDP-43 target[4–6] | -8.439 | -9.004 |
| ***PTGDS*** | Prostaglandin D2 synthase | Functions as a neuromodulator and trophic factor in the CNS; known TDP-43 target[4] | -3.579 | -3.477 |
| *ZNF717* | Zinc finger protein 717 | Kruppel-associated box (KRAB) zinc-finger protein; transcriptional regulator | -7.149 | -9.269 |
| *LINC02506* | Long intergenic non-protein coding RNA 2506 | lncRNA | -6.348 | -4.170 |
| ***ZNF283*** | Zinc finger protein 283 | Predicted to enable DNA-binding transcription factor activity | -3.759 | -3.404 |
| *SVIL-AS1* | SVIL antisense RNA 1 | lncRNA | -7.275 | -7.288 |
| ***RAD51C*** | RAD51 paralog C | Homologous recombination and DNA repair | -0.455 | -0.301 |
| ***PCDHA13*** | Protocadherin alpha 13 | Neural cadherin-like cell adhesion protein; enriched in the brain | -6.492 | -6.244 |
| *ZNF736* | Zinc finger protein 736 | Predicted to enable DNA-binding transcription factor activity | -5.283 | -6.767 |
| *ENSG00000229370* | Novel transcript | lncRNA | -8.090 | -6.443 |
| *ENSG00000287069* | Novel transcript, antisense to *KCNH8* | lncRNA | -4.777 | -4.134 |
| ***NNAT*** | Neuronatin | May play a role in the formation and maintenance of the CNS, and regulation of ion channels during brain development | -10.147 | -11.453 |
| ***GPR50*** | G protein-coupled receptor 50 | G-coupled receptor activity; associated with bipolar affective disorder and major depressive disorder | -3.561 | -3.455 |
| ***CHCHD2*** | Coiled-coil-helix-coiled-coil-helix domain containing 2 | Involved in mitochondrion organization; negative regulator of mitochondria-mediated apoptosis; associated with PD; predicted ALS gene[3] | -10.399 | -11.938 |

| | | | | |
|---|---|---|---|---|
| *LINC02527* | Long intergenic non-protein coding RNA 2527 | lncRNA | -0.873 | -1.342 |
| *MIR4458HG* | MIR4458 host gene | lncRNA | -10.030 | -9.931 |
| *CCNYL2* | Cyclin Y like 2 | Predicted to enable cyclin-dependent protein serine/threonine kinase regulator activity | -4.539 | -4.344 |
| ***PDE12*** | Phosphodiesterase 12 | Localized to the mitochondrial matrix; involved in RNA metabolic process, cytokine signaling and mitochondrial mRNA stability | -0.392 | -0.393 |
| ***CBR1*** | Carbonyl reductase 1 | NADPH-dependent oxidoreductase activity; linked to fatty acid metabolism | -0.834 | -0.615 |
| *ABCA4* | ATP binding cassette subfamily A member 4 | Retina-specific ATP-binding cassette (ABC) transporter; associated with retinal diseases | -2.794 | -2.913 |
| ***Discordant*** | | | | |
| *ASNSP1* | Asparagine synthetase pseudogene | Pseudogene | 2.776 | -5.693 |
| *MYH14* | Myosin heavy chain 14 | Non-muscle myosin involved in cytokinesis, cell motility, and cell polarity; associated with hearing impairment and peripheral neuropathy; predicted ALS gene[3] | -5.120 | 4.971 |
| *NBEAP1* | Neurobeachin pseudogene 1 | Pseudogene | 1.213 | -5.991 |
| *BMS1P10* | BMS1 pseudogene 10 | Pseudogene | 1.785 | -2.262 |
| *ENSG00000272234* | Novel transcript, antisense to *SEPP1* | lncRNA | 2.231 | -3.091 |
| *CCDC152* | Coiled-coil domain containing 152 | Protein-coding gene of unknown function | 2.421 | -4.929 |
| *PUS7L* | pseudouridine synthase 7 like | Predicted to enable pseudouridine synthase activity; associated with epilepsy | 0.333 | -0.610 |
| *ZSCAN1* | Zinc finger and SCAN domain containing 1 | DNA-binding transcription factor activity | 1.697 | -9.783 |

* ALS, amyotrophic lateral sclerosis; CNS, central nervous system; FC, fold change; lncRNA, long non-coding RNA; PD, Parkinson's disease

** Bolded genes selected for validation by qPCR

*** According to the GeneCards database available at https://www.genecards.org/

**Table S6. Shared dysregulated miRNAs and their genomic location**

| miRNA | Genomic coordinate | | LogFC p.A382T | LogFC p.G348C |
|---|---|---|---|---|
| hsa-miR-6832-3p | chr6: 31633787-31633858 [+] | 6p21.33 | 5.39753821 | 5.09015244 |
| hsa-miR-377-5p | chr14: 101062050-101062118 [+] | 14q32.31 | -3.4987295 | -3.5640906 |
| hsa-miR-432-5p | chr14: 100884483-100884576 [+] | 14q32.2 | -8.5096634 | -7.6425164 |
| hsa-miR-370-3p | chr14: 100911139-100911213 [+] | 14q32.31 | -6.1090404 | -5.9600407 |
| hsa-miR-485-3p | chr14: 101055419-101055491 [+] | 14q32.31 | -2.7175301 | -3.2348732 |
| hsa-miR-487a-3p | chr14: 101052446-101052525 [+] | 14q32.31 | -2.9450634 | -3.1297286 |
| hsa-miR-134-5p | chr14: 101054687-101054759 [+] | 14q32.31 | -2.7123204 | -2.9936839 |
| hsa-miR-495-3p | chr14: 101033755-101033836 [+] | 14q32.31 | -2.8734665 | -2.7943359 |
| hsa-miR-329-3p | chr14: 101026785-101026864 [+]<br>chr14: 101027100-101027183 [+] | 14q32.31 | -2.7681077 | -3.276883 |
| hsa-miR-758-3p | chr14: 101026020-101026107 [+] | 14q32.31 | -3.1996719 | -3.1022029 |
| hsa-miR-487b-3p | chr14: 101046455-101046538 [+] | 14q32.31 | -3.191169 | -3.2420759 |
| hsa-miR-654-5p | chr14: 101040219-101040299 [+] | 14q32.31 | -2.8213098 | -2.9534777 |
| hsa-miR-412-5p | chr14: 101065447-101065537 [+] | 14q32.31 | -2.9474603 | -3.6345566 |
| hsa-miR-656-3p | chr14: 101066724-101066801 [+] | 14q32.31 | -2.9487844 | -4.1468312 |
| hsa-miR-323a-3p | chr14: 101025732-101025817 [+] | 14q32.31 | -2.6949508 | -2.9099091 |
| hsa-miR-376a-3p | chr14: 101040782-101040849 [+]<br>chr14: 101040069-101040148 [+] | 14q32.31 | -3.4993597 | -3.7251577 |
| hsa-miR-409-3p | chr14: 101065300-101065378 [+] | 14q32.31 | -3.2102507 | -3.0777423 |
| hsa-miR-381-3p | chr14: 101045920-101045994 [+] | 14q32.31 | -3.132852 | -3.4426206 |
| hsa-miR-889-3p | chr14: 101047901-101047979 [+] | 14q32.31 | -3.040022 | -3.4464336 |
| hsa-miR-485-5p | chr14: 101055419-101055491 [+] | 14q32.31 | -2.8677297 | -3.1330652 |
| hsa-miR-136-3p | chr14: 100884702-100884783 [+] | 14q32.2 | -7.2324032 | -6.4719432 |
| hsa-miR-411-5p | chr14: 101023325-101023420 [+] | 14q32.31 | -2.8392333 | -3.491725 |
| hsa-miR-433-3p | chr14: 100881886-100881978 [+] | 14q32.2 | -6.6722867 | -6.9097905 |
| hsa-miR-379-3p | chr14: 101022066-101022132 [+] | 14q32.31 | -3.6183613 | -2.809508 |
| hsa-miR-382-5p | chr14: 101054306-101054381 [+] | 14q32.31 | -2.9233342 | -3.2294855 |
| hsa-miR-411-3p | chr14: 101023325-101023420 [+] | 14q32.31 | -3.1726077 | -3.1718531 |
| hsa-miR-493-5p | chr14: 100869060-100869148 [+] | 14q32.2 | -8.8452075 | -6.2919642 |
| hsa-miR-369-5p | chr14: 101065598-101065667 [+] | 14q32.31 | -3.348575 | -3.6018686 |
| hsa-miR-379-5p | chr14: 101022066-101022132 [+] | 14q32.31 | -2.9522166 | -3.1315874 |
| hsa-miR-493-3p | chr14: 101033755-101033836 [+] | 14q32.31 | -7.4756617 | -5.4134923 |
| hsa-miR-1185-2-3p | chr14: 101044198-101044283 [+] | 14q32.31 | -3.5779433 | -3.6520981 |
| hsa-miR-127-3p | chr14: 100882979-100883075 [+] | 14q32.2 | -6.2099569 | -6.7713807 |
| hsa-miR-654-3p | chr14: 101040219-101040299 [+] | 14q32.31 | -2.9824453 | -3.1408796 |
| hsa-miR-655-3p | chr14: 101049550-101049646 [+] | 14q32.31 | -3.1648455 | -3.6508573 |
| hsa-miR-543 | chr14: 101031987-101032064 [+] | 14q32.31 | -2.9567885 | -2.6150719 |
| hsa-miR-410-3p | chr14: 101065912-101065991 [+] | 14q32.31 | -3.0472873 | -3.2986066 |

* FC, fold change

**Table S7. Predicted TDP-43 binding sites by the RBPmap database**[7]

| Gene | Genomic coordinate | Motif | Predicted binding sites | Z-score | *P*-value |
|---|---|---|---|---|---|
| *MIR377* | chr14:101062114 | guaug | guugaaucacacaaaggcaacuuuuguuug | 2.214 | 1.34E-02 |
| *MIR370* | chr14:100911182 | guaug | acgucucugcaguuacacagcucacgagugccugcuggggguggaaccuggucugu | 2.051 | 2.01E-02 |
| | chr14:100911194 | guaug | uuacacagcucacgagugccugcugggguggaaccuggucugucu | 2.316 | 1.03E-02 |
| | chr14:100911206 | guaug | cgagugccugcuggggguggaaccuggucugucu | 2.714 | 3.32E-03 |
| *MIR485* | chr14:101055424 | gagug | acuuggagagaggcuggccgugaugaauucgauuc | 2.073 | 1.91E-02 |
| | chr14:101055426 | gagug | acuuggagagaggcuggccgugaugaauucgauucau | 2.073 | 1.91E-02 |
| *MIR134* | chr14:101054689 | guaug | cagggugugugugacugguugaccagaggggcau | 2.469 | 6.77E-03 |
| | chr14:101054691 | guaug | caggguguguugacugguugaccagaggggcaugc | 3.194 | 7.02E-04 |
| | chr14:101054693 | guaug | cagggugugugacugguugaccagaggggcaugcac | 3.194 | 7.02E-04 |
| | chr14:101054697 | guaug | cagggugugugacugguugaccagaggggcaugcacugug | 2.551 | 5.37E-03 |
| | chr14:101054710 | gagug | agggugugugacugguugaccagagggggcaugcacuguguucacccuguggg | 2.303 | 1.06E-02 |
| | chr14:101054715 | guaug | ggugugugacugguugaccagagggggcaugcacuguguucacccugugggccacc | 2.918 | 1.76E-03 |
| *MIR654* | chr14:101040221 | guaug | ggguaaguggaaagaugguggggccgcagaaca | 1.684 | 4.61E-02 |
| *MIR412* | chr14:101065458 | guaug | cugggguacggggaugggauggucgaccaguuggaaaguaau | 1.776 | 3.79E-02 |
| | chr14:101065462 | guaug | cugggguacggggauggauggucgaccaguuggaaaguaauuguu | 1.776 | 3.79E-02 |
| | chr14:101065525 | guaug | acuucaccugguccacuagccguccguauccgcugcag | 1.673 | 4.72E-02 |
| *MIR656* | chr14:101066758 | guaug | guugccugugagguguucacuuucuauaugaugaauauuauacagucaaccucuu | 1.684 | 4.61E-02 |
| *MIR376A2* | chr14:101040094 | guaug | gguauuuaaaagguagauuuuccuucuaugguuacguguuugaugguuaaucaua | 1.929 | 2.69E-02 |
| *MIR889* | chr14:101047910 | guaug | gugcuuaaagaauggcuguccguaguauggucucuauau | 2.306 | 1.06E-02 |
| | chr14:101047925 | guaug | gugcuuaaagaauggcuguccguaguaugggucucuauauuuaugaugauuaaua | 3.041 | 1.18E-03 |
| *MIR411* | chr14:101023349 | guaug | ugguacuuggagagauaguagaccguauagcguacgcuuuaucugugacguaug | 2.276 | 1.14E-02 |
| | chr14:101023356 | guaug | uuggagagauaguagaccguauagcguacgcuuuaucugugacguauguaacacg | 2.276 | 1.14E-02 |
| | chr14:101023374 | guaug | guauagcguacgcuuuaucugugacguauguaacacgguccacuaacccucagua | 2.786 | 2.67E-03 |
| *MIR379* | chr14:101022087 | guaug | agagaugguagacuauggaacguaggcguuaugauuucugaccuauguaac | 1.816 | 3.47E-02 |
| | chr14:101022094 | guaug | gaugguagacuauggaacguaggcguuaugauuucugaccuauguaacauggucc | 1.816 | 3.47E-02 |
| *MIR493* | chr14:100869128 | guaug | uuugcacauucggugaaggucuacugugugccaggcccugugccag | 1.755 | 3.96E-02 |

**Table S9. Top-scoring predicted compounds by the CMap database**[8]

| Name | Description | τ score | | |
|---|---|---|---|---|
| | | p.A382T | p.G348C | Combined |
| sulpiride | Dopamine receptor antagonist | 0.00 | 1.35 | -94.56 |
| tacedinaline (CI-994) | HDAC inhibitor | -1.02 | -0.92 | -92.30 |
| XMD-892 | MAP kinase inhibitor | 0.00 | 11.26 | -91.09 |
| QS-11 | ARFGAP inhibitor | -0.49 | -14.82 | -90.83 |
| bimatoprost | Prostanoid receptor agonist | 0.00 | 0.00 | -90.58 |
| XMD-885 | Leucine rich repeat kinase inhibitor | -40.03 | -5.53 | -90.49 |
| GR-46611 | Serotonin receptor agonist | -1.30 | -2.04 | -90.17 |
| CS-110266 | Dopamine receptor agonist | -1.21 | 14.28 | -89.57 |
| benzohydroxamic-acid | Antifungal | 1.80 | -2.33 | -88.57 |
| MLN4924 (Pevonedistat) | Nedd activating enzyme inhibitor | 0.39 | -49.65 | -87.49 |
| piperlongumine | Glutathione transferase inhibitor | -71.64 | 1.06 | -87.05 |
| alpha-linolenic-acid | Omega 3 fatty acid stimulant | 0.00 | 1.62 | -85.92 |
| 4-hydroxy-2-nonenal | Cytotoxic lipid peroxidation product | 0.00 | 0.00 | -85.32 |
| prunetin | Breast cancer resistance protein inhibitor | -95.71 | 29.48 | -71.45 |
| mitomycin-c | DNA alkylating agent | -93.55 | 74.75 | -69.90 |
| zaldaride | Calmodulin antagonist | 0.00 | -88.60 | 0.60 |
